# Representation of grasping type and force in the primate motor cortex

**DOI:** 10.1101/2024.05.23.595565

**Authors:** Adriana Moreno, Victor de Lafuente, Hugo Merchant

**Author notes:** **Corresponding authors:** Hugo Merchant and Victor de Lafuente. The authors declare no competing financial interests.

## Abstract

The primary motor cortex (M1) is strongly engaged by movement planning and movement execution. However, the role of M1 activity in voluntary grasping is still not completely understood. Here we analyze recordings of M1 neurons during the execution of a delayed reach-to-grasp task, where monkeys had to actively grasp an object with either a side or a precision grip, and then pull it with a low or high amount of force. Single cell and neural populations analyses showed that grip type was robustly and specifically encoded by a large population of neurons, while force level was weakly encoded within mixed-selective neurons that also provided grip type information. Notably, the grip type was stably decoded from motor cortical populations during the preparation and execution epochs of the task. Our results are consistent with the idea that planning and performing specific grasping movements are high-level skills that strongly engage M1 neurons, while the execution of grasping-pulling force might be prominently encoded at lower stages of the motor system.

**Significance statement:** Grasping behavior requires precise motor coordination exerted by multiple brain areas, including the primary motor cortex (M1), but the exact role of M1 in grasping preparation and execution remains elusive. Here, we analyzed the neural activity of M1 while two monkeys performed a delayed reach-to-grasp task. We found that two parameters of grasping: grip type and force level, were represented in the activity of single neurons and the neural population, although with important differences. While grip representation was stronger and more temporally stable, force encoding was weaker and short lived. Our results suggest that grip planning and execution is a high-level neural process that takes place independently of force control in M1.

## Introduction

Primates, including humans, often manipulate objects using their sophisticated prehensile hands. After determining an object’s position in space, we must estimate object features like size, shape, texture, and weight. This information is then used by the brain to plan and execute precise movements to hold it appropriately. These movements comprise two main phases: a reaching phase in which the hand approaches the object, and a grasping phase that occurs once contact is made. During the reaching phase, as the hand moves, the position of the fingers changes, adapting to features of the object we intend to grasp (*preshaping*). During the grasping phase, the position of the hand keeps adapting as the brain now receives feedback from tactile and proprioceptive inputs (Jeannerod et al., 1995). Human and monkey hands are capable of different types of grips, including the highly controlled precision grip, in which an object is held between the palmar aspects of the fingers and the opposing thumb (Almécija & Sherwood, 2016). Besides controlling hand position and its derivatives, it is also crucial to regulate the amount of grasping force. If we apply too little force, the object may slip out of our hands, whereas if we apply too much, it may break or become damaged. Throughout the years, efforts have been made to determine how the brain gives rise to this highly precise and controlled behavior.

Due to M1’s spinal projections, earlier studies aimed to find correlations between M1 neural activity and muscle activation. Surprisingly, the responses of M1 neurons showed to be highly diverse. While the firing rates of some neurons do correlate with muscle activation during single-joint movements and isometric force control (Cheney & Fetz, 1980; Evarts, 1968; Maier et al., 1993; Wannier et al., 1991), others seem to carry information about parametric aspects of kinematics like direction, speed, trajectory and dynamic force changes (Georgopoulos et al., 1986, 1992, 2007; Hatsopoulos et al., 2007; Hepp-Reymond et al., 1999; Inoue et al., 2018). In addition, recent studies have revealed that M1 neural populations can be described as a dynamical system in which internally generated temporal patterns generate motor commands that are sent to the spinal motoneurons and eventually, effector muscles (Churchland et al., 2012; Russo et al., 2018).

In addition to movement execution, previous studies have shown that M1 activity correlates with movement preparation, with population responses that converge to a particular state space when the system has encoded all the properties of the upcoming reaching movement (Afshar et al., 2011; Churchland et al., 2006; Riehle & Requin, 1993). A recent paper demonstrated that this principle also generalizes for grasping movements. It was shown that M1 preparatory activity encodes different types of grips with different locations in the population state space. In addition, when monkeys prepare two potential grips in parallel, the activity of M1 situates within an intermediate preparatory state, which constitutes an optimal initial condition to promptly execute either of the two planned movements (Meirhaeghe et al., 2023).

Here we investigated how M1 neurons represent two crucial aspects of grasping: grip type and grip force. We performed encoding and decoding analyses on the activity of simultaneously recorded M1 neurons from two monkeys performing a delayed reach-to-grasp task (Brochier et al., 2018). We found that at the single-unit and population levels, the type of grip had a profound effect on the motor cortical activity while force level explained less variance of the neural responses. This was achieved through a combination of both more neurons representing grip, as well as enhanced grip coding at the single-unit level.

Notably, several neurons were modulated only by grip type, while force was represented by neurons selective for both grip and force. Furthermore, the neural signals linked to grip type emerged right after grip instruction and were maintained after movement onset, forming a long-lasting and stable preparatory signal for precision or side grip that extended through movement execution. Importantly, even while pulling the object, when force level and grip type are simultaneously being executed, force level engaged fewer number of neurons and accounted for a small proportion of activity variance.

## Materials and methods

### Behavioral task

We used the online database generously made public by Brochier et. al (2018). The task and recording methods are explained in detailed elsewhere (Brochier et al., 2018). Briefly, two macaque monkeys (N and L) were trained in an instructed delayed reach-to-grasp task, in which animals were visually instructed to grasp an object using either a side grip or a precision grip. After grasping, the monkeys had to pull the object for at least 500 ms with either a low or a high force level. If the monkeys followed the instructions correctly, they were rewarded with a drop of apple sauce at the end of the trial. Figure 1 shows the sequence of events that defined each trial of the grasping task. Note that grip type and pulling-force cues were temporally separated. The monkeys were first instructed about grip type, and after a 1 s delay, they were given an additional visual cue to indicate force magnitude. The force level cue also served as the go-cue to initiate the hand movement. Thus, monkeys had more time to prepare the type of grip, and less time to plan the grip force required to pull the object. Reaction time (RT) was defined as the elapsed time between the go-cue and the start of the reaching movement. Movement time (MT) was calculated as the time from movement onset until the object was touched.

**Figure 1.**
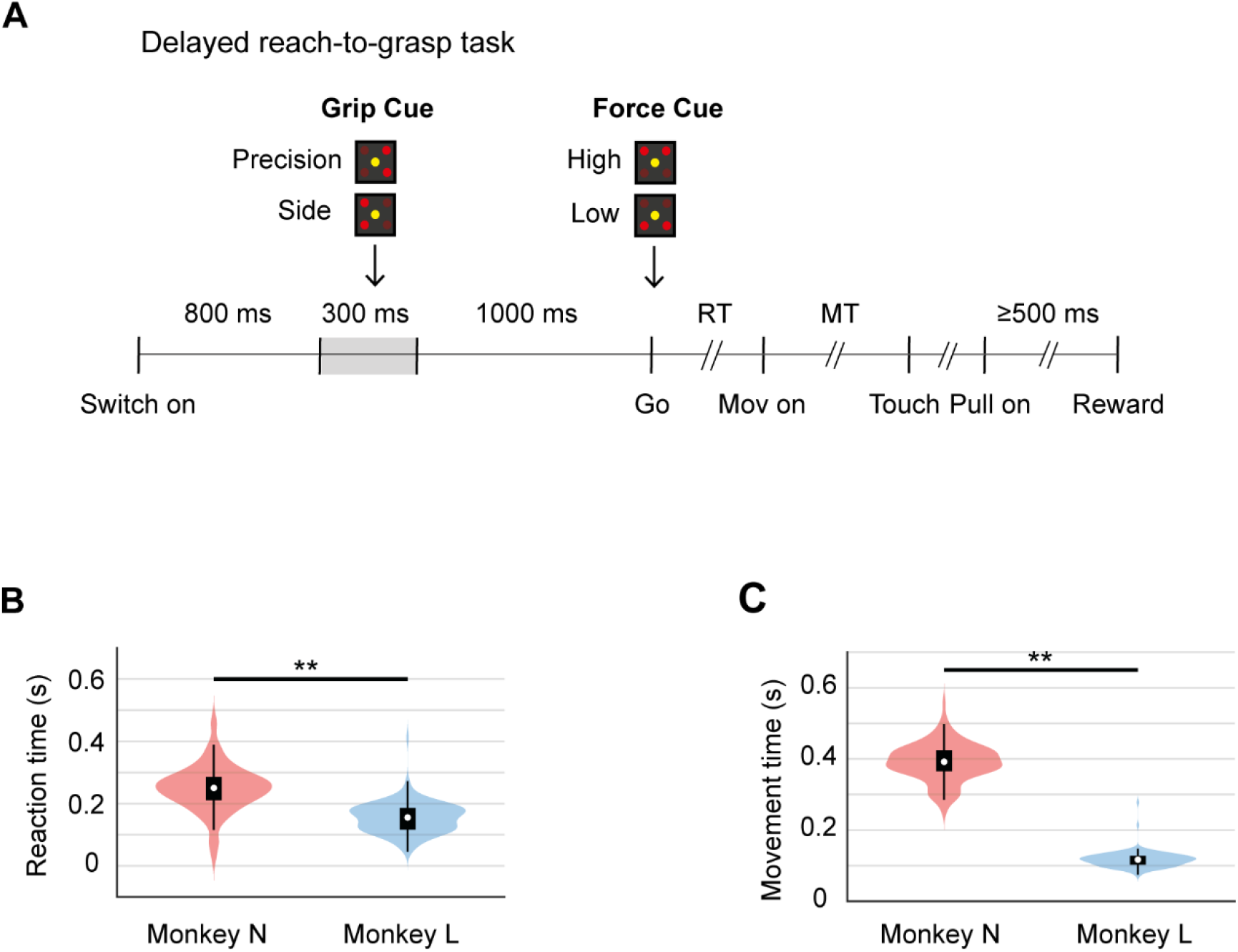
Delayed reach-to-grasp task. A: In this task, monkeys are instructed to grasp and pull an object using a specific hand grip and force level. Each trial starts with the monkey placing its working hand over a table switch (Switch On). After 800 ms, a visual cue is given to the monkey to indicate the hand grip (a precision grip or a side grip) he must use to grasp the object. After a 1000 ms delay, another visual cue (Go) instructs the monkey to start moving, as well as the amount of force (low or high) needed to pull the object. Reaction time (RT) was measured as the time between the Go cue and the movement onset (Movement On). Movement time (MT) was defined as the time between the movement onset and the object touch (Touch). Once grasped, the monkey must pull the object (Pull On) for at least 500 ms with the instructed force level. If the monkey performs correctly, he is rewarded with a drop of apple sauce (Reward). B: Reaction times during correct trials for monkey N and monkey L, respectively. C: Movement times for both monkeys, displayed as in B.

### Neural recordings

A 10-by-10 Utah electrode array (Blackrock Microsystems, Salt Lake City, UT, USA) was chronically implanted in the primary motor (M1) and premotor cortex (dorsally in monkey L and more ventrally in monkey N) on the right hemisphere of each monkey, contralateral to the working hand. Neural signals coming from a total of 96 electrodes, were amplified, high pass filtered (0.3 Hz – 7.5 kHz) and digitized with a sampling rate of 30 kHz. Raw signals were online high-pass filtered at 250 Hz and spike waveforms were detected by threshold crossing (manually set). To isolate single-unit activity, both online and offline sorting were performed (Plexon Offline Spike Sorter, Plexon Inc, Dallas, Texas, USA). We focused only on motor cortical activity, using neural signals from electrodes located posterior to the putative border between M1 and the premotor cortex (a total of 61 sites for monkey N and 68 sites for monkey L).

### Single-unit analysis

To study activity at single-unit level, we calculated the firing rate of each neuron as a function of time by using an exponential filter with a decay rate of 100 ms displaced every 20 ms. Normalized activity (z-score) was calculated for each neuron by subtracting its mean firing rate (across trials and time) to the firing rate of every time point in every trial and then dividing it by its standard deviation. For each neuron, firing rates were averaged across trials and aligned to the onset of the reaching movement.

To determine whether the firing rate of individual neurons contained information about the task’s variables, we calculated mutual information for grip type and force level for each neuron as a function of time. The amount of information carried by the firing rate of a neuron *N* about a task parameter *x* can be estimated as:

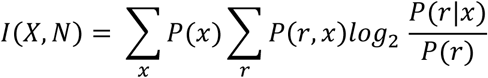

Where *P(x)* is the probability of task parameter *x, P(r)* corresponds to the probability of occurrence of neuronal response r among all possible responses and *P(r*|*x)* is the conditional probability of obtaining a neuronal response *r* given the presence of the task parameter *x*. To identify neurons that carried a significant amount of mutual information about grip type or force level, we performed a non-parametric permutation test. For each neuron, we shuffled its firing rate values across trials and time, and calculated the permuted mutual information as a function of time. We repeated this procedure 1000 times and considered the neuron’s amount of mutual information to be significant when the mutual information values were above chance (p<0.05) for at least 8 consecutive bins.

To characterize firing rate modulations of individual neurons in response to grip and force conditions, we calculated the receiver operating characteristic (ROC) index as a function of time. The ROC index estimates the overlap between two distributions. For each neuron, the distributions compared the firing rates of one condition versus its counterpart condition (side grip trials vs precision grip trials; and high force trials vs low force trials). A ROC value of 0.5 indicates that distributions completely overlapped and thus, firing rates are not useful to differentiate the two conditions. We calculated each neuron’s ROC index as a function of time and considered the ROC values that were significantly different from 0.5 (95% confidence interval). If ROC values were significant for at least 10 bins, we considered that neuron to have a significant ROC index and took the value when the absolute difference between 0.5 was maximum (de Lafuente & Romo, 2006; Merchant et al., 1997).

### Neural population analysis

To determine whether M1 encodes grip type and force level at the neural population level, we performed a demixed principal component analysis (Kobak et al., 2016). Similarly to principal component analysis (PCA), dPCA is a dimensionality-reduction method that allows to extract the neural population components that explain most variance (Gámez et al., 2019). In dPCA these components are referred to as ‘demixed’ because they explain variance associated with specific task variables. In this case, the demixed principal components (dPCs) were associated with grip type, force level, and temporal events within the task (referred to as condition-independent activity). Additionally, to detect the time points at which task parameters were significantly encoded by the neural population at single-trial level, significant dPCs were used as linear classifiers. For each dPC, a stratified Monte Carlo leave-group-out cross-validation was performed. Monte Carlo chance distribution was set to 100 and time periods when the actual classification accuracy exceeded shuffled decoding accuracies in at least 10 consecutive time bins were considered significant.

### Neural population decoding

A decoding analysis was implemented using a methodology developed by Crowe et al., (Crowe et al., 2014), and Merchant and Averbeck (Merchant & Averbeck, 2017). For each neuron, the discharge rate was computed in 100 ms time bins within -2000 to 2000 ms relative to movement onset, resulting in 51 time-bins, across the 30 trials of the two grip and two force conditions. For each time bin, we built an SVM classifier with a 10-fold crossvalidation. The procedure was repeated 50 times using different training (n = 25) and testing (n = 5) trials each time, and the prediction results were averaged between repetitions. Accuracy thus estimated classifier’s ability to discriminate the experimental conditions and corresponds to the total of correctly predicted trials divided by the total number tested conditions (2 for grip type and 2 for force level) by the number of toral trials. A 50% accuracy corresponds to a random classification performance for the two levels of the parameters.

We carried out a cross-temporal decoding analysis in which we trained and tested the SVMs using all possible combinations of 100-ms time bins to establish the temporal evolution of information coding. This analysis results in a classification accuracy matrix where the values along the diagonal are calculated by performing training and testing on equivalent time bins (Figures 7B, 8B). In contrast, different time bins are used for training and testing and calculating the off-diagonal accuracy values. The off-diagonal accuracy allows us to determine how stable in time is a neural population signal when compared with the on-diagonal values. In fact, we considered an off-diagonal accuracy bin-pair combination as static, namely, a bin combination that shows decoding generalization from the on-diagonal bin classification model, when the three following conditions were met. The offdiagonal accuracy was significantly higher than chance (cluster-based permutation test, p < 0.01), was above the 99% of the bootstrapped null distribution (1000 iterations with shuffled classification labels), and that the two corresponding time bins on-diagonal showed a classification accuracy above chance level (permutation test, p < 0.01, Bonferroni corrected for the number of on-diagonal time bins). Next, we computed a binary matrix of the same size as the cross-temporal decoding matrix, where we assigned 1 to at least two consecutive time bins resulting as static and 0 for all remaining time bins. Then, we quantified the magnitude of cross-time decoding between and within the two main task epochs using the generalization index. The generalization index provides information about the proportion of SVM tested bins during the preparation and execution that were classified as static during the trained bins of either the preparation or execution epoch.

## Results

### Monkeys were able to use two grip types and two force levels to grasp and pull an object

In the delayed reach-to-grasp task monkeys were instructed to grasp a cubic object using either a side or a precision grip. Once grasped, the monkeys had to pull and hold the object for at least 500 ms with a low or high amount of force. Figure 1A shows the main events of the task. Each trial started with the monkey placing its left hand over a table switch. After 800 ms, a visual cue was displayed for 300 ms, instructing the monkey about grip type. After a 1000 ms delay, an additional visual cue indicated the required force level to pull the object. This cue also served as a go cue for the monkey to start the reaching movement. Thus, this task was designed to determine the neural underpinnings of movement preparation and execution of multijoin complex hand movements, and the neural signals related with the execution of two types of grasping movements combined with two levels of grasping force.

The behavioral results show that monkey L performed the task faster, with shorter and less variable reaction times (Figure 1B; mean ± SD: 150 ms ± 48 ms) compared to monkey N (mean ± SD: 251 ms ± 190 ms). Movement times were also shorter and less variable in monkey L as compared to monkey N (Figure 1C; mean ± SD; monkey L: 123 ms ± 80 ms, monkey N: 343 ms ± 125 ms). Both parameters were significantly different between animals (reaction time: z = -11.51, p = 1.07×10^-30^; movement time: z = -14.01, p = 1.29×10^-44^, Mann Whitney U-test).

### Grip type has a larger influence in M1 neurons as compared to grip force

While the monkeys performed the reach-to-grap task, extracellular spike potentials from M1 neurons were recorded. Figure 2A shows the raster plot and firing rate of a neuron from monkey N that increased its activity after movement onset, with a preference for side over precision grip trials. Note that, as object touch approximates, activity started to become modulated also by force level. Figure 2B shows a neuron that fired strongly in precision grip trials. Figure 2C illustrates a neuron that fired strongly during the preparation and execution of the side grip in low force trials. Finally, Figure 2D depicts a neuron that is more active during preparation and execution of a precision grip and preferred high force trials once the monkey touched the object.

**Figure 2.**
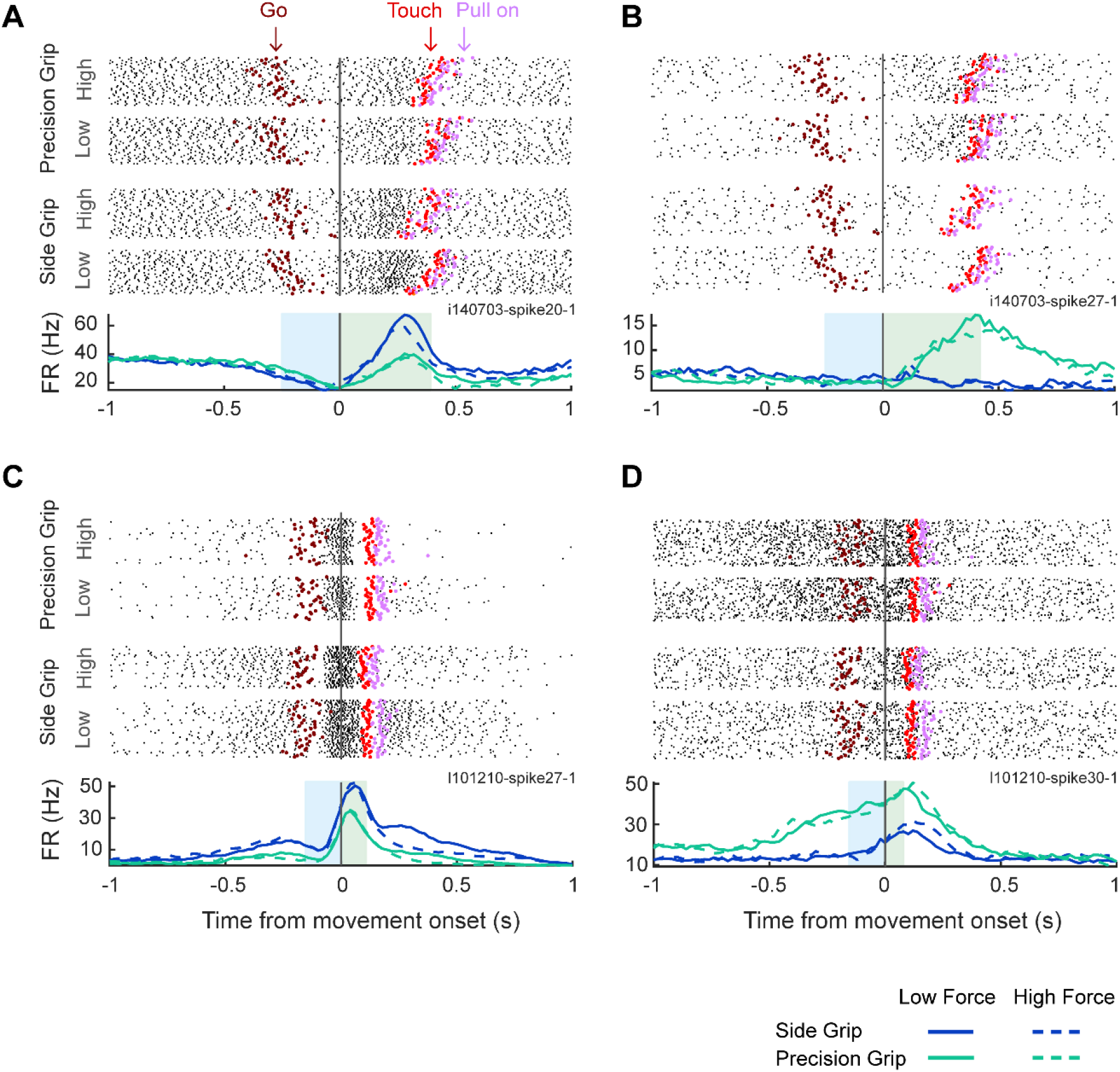
Four example neurons of M1 during the delayed reach-to-grasp task. A-B (top): Raster plot of two example neurons from monkey N. Trials are grouped according to the instructed grip type and force level, as indicated by the legends on the left (from bottom to top: side grip and low force, side grip and high force, precision grip and low force, precision grip and high force). Activity is aligned to the movement onset (grey solid line). Colored markers indicate task events around the movement onset: Go cue (dark red), object touch (red), pull onset (lilac). A-B (bottom): Trial-averaged firing rate across task conditions. Line color indicates grip type (blue for side grip and green for precision grip). Line style indicates force level (dashed line for high force and solid line for low force). Blue shading indicates mean reaction time and green shading indicates mean movement time. C-D: Activity of two example neurons from monkey L as shown in A and B.

To determine whether single-unit activity contained information about task parameters, we computed the amount of mutual information associated with grip type and force level, and we did this for each neuron, before and after movement onset using a sliding window (see Methods). The heatmaps in Figure 3A-B show mutual information as a function of time, for neurons with significant effects for either grip type (left panel) or force (right panel; permutation test, p < 0.001). Neurons were sorted by the time of their maximum mutual information value. In monkey N, the mutual information peaked after movement onset for most neurons, while maximum force information occurred later after object touch.

**Figure 3.**
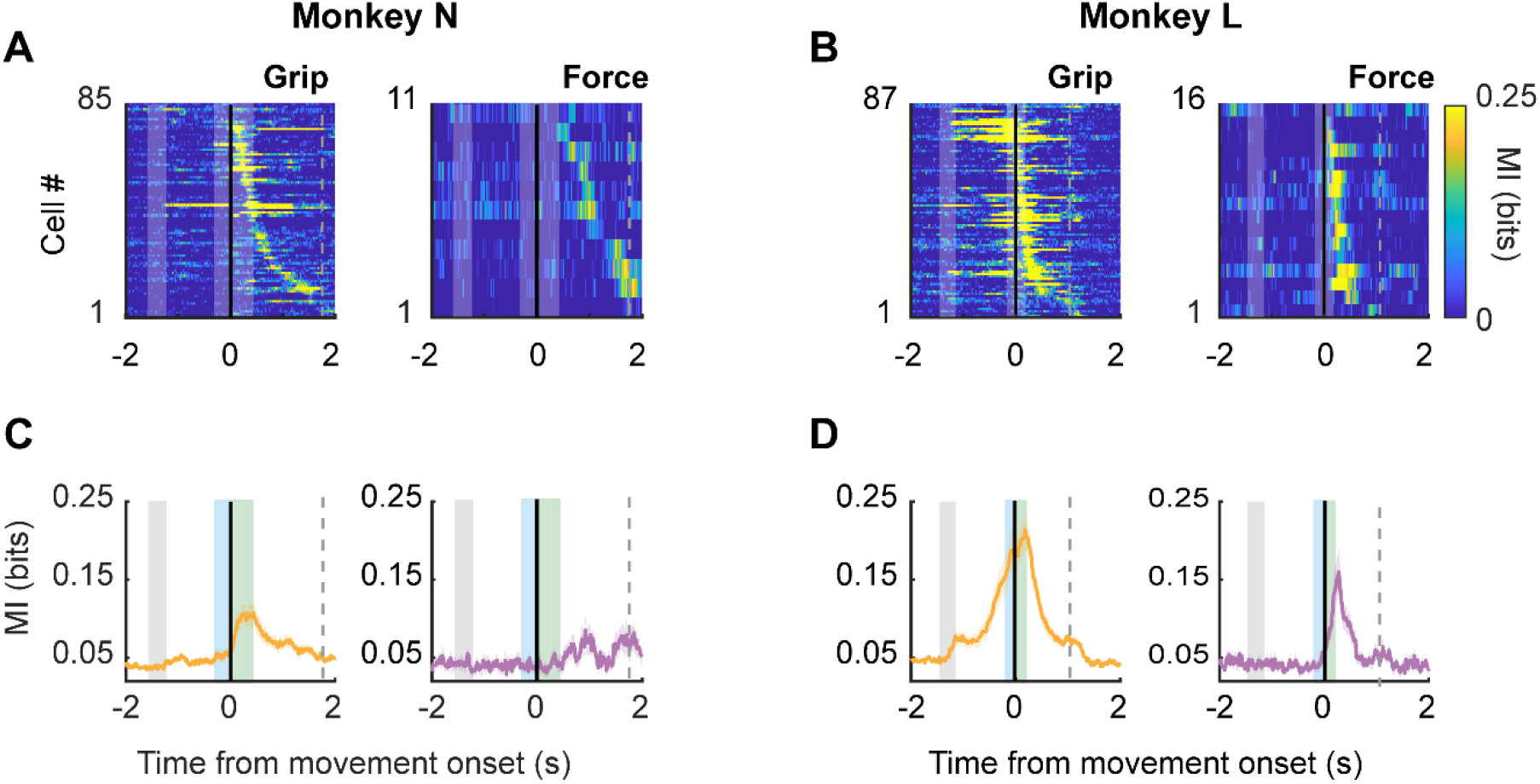
Grip and force-related mutual information of M1 neurons during the task. A: Heatmaps of grip (left) and force (right) mutual information for monkey N. Only neurons that carry a significant amount of mutual information (p<0.01) are shown. B: Same as in A, for monkey L. C: Average grip (left) and force (right) mutual information across neurons as a function of time for monkey N. Grey shading indicates the duration of the grip cue. Blue and green shading indicate mean reaction and movement times, respectively. Black solid line indicates movement onset. Grey dashed line indicates mean object release time. D: Same as in C, for monkey L.

The time profile of mutual information clearly indicated that neurons carried less information about force in comparison to grip (Figure 3C-D). Note that monkey L had larger and earlier peaking mutual information of both parameters compared to monkey N, with a strong preparatory signal for grip type.

Next, we assessed whether grip and force were encoded by different or shared neural subpopulations by labeling neurons according to the parameter they significantly carried information about. Panels A and B of Figure 4 show the average mutual information of neurons that carried both grip and force information for monkey N (n = 10 neurons) and monkey L (n = 16 neurons), respectively. Figure 4C and 4D show the mean mutual information of neurons that carried only grip information in monkey N (n = 75 neurons) and monkey L (n = 71).

**Figure 4.**
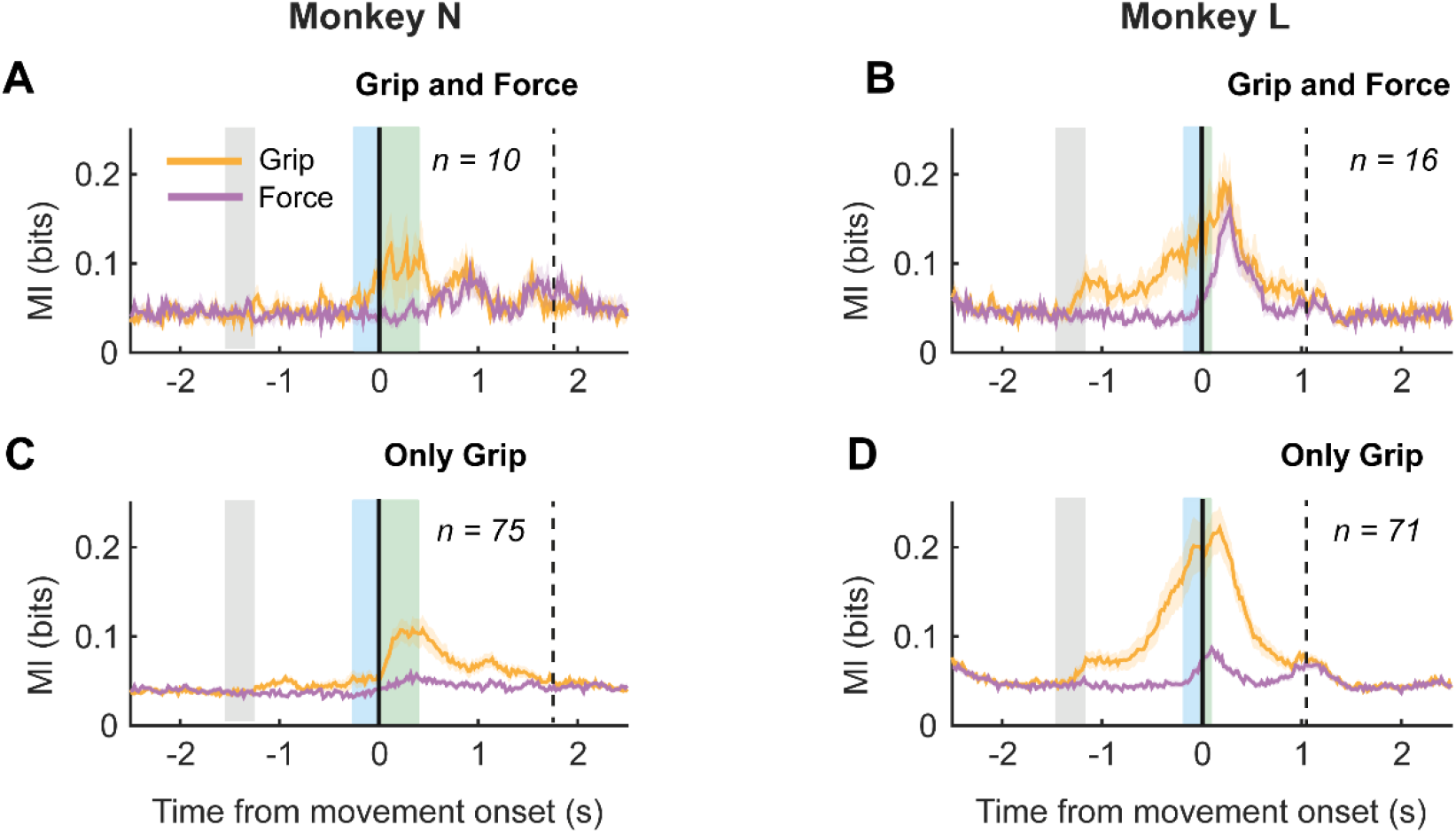
Force is represented in M1 by a mixed-selective subpopulation for both grip and force. A: Average mutual information related to grip (yellow solid line) and force (purple solid line) as a function of time for neurons that carry grip and force information in monkey N (n = 10 neurons). Grey shading indicates the duration of the grip cue. Blue and green shading indicate mean reaction and movement times, respectively. Black solid line indicates movement onset. B: Same as in A, for monkey L (n = 16 neurons). C: Average mutual information for neurons carrying significant grip information only, for monkey N (n = 75 neurons). D: Same as in C, for monkey L (n = 71 neurons).

Notably, no subpopulation contained force-only information, and this was true for both monkeys. Therefore, the prevalent grip type signal mainly depended on a large population of grip specific cells that were active during the preparation and execution epochs. Conversely, force was represented by a smaller neural subpopulation that also contained grip representations during movement execution.

Once we identified which neural subpopulations contained grip and force representations, we sought to determine if grip and force were represented similarly in the firing rate of individual units and if such neural codes were dependent of each other. To achieve this, we estimated the ROC index comparing neural activity during both grip types and force levels for all neurons during the movement preparation and execution epochs (Methods). Figure 5A-B shows the maximum ROC value of the comparison between side grip vs precision grip trials in the y-axis with the maximum ROC value of the comparison between high vs low force in the x-axis (see Methods).

**Figure 5.**
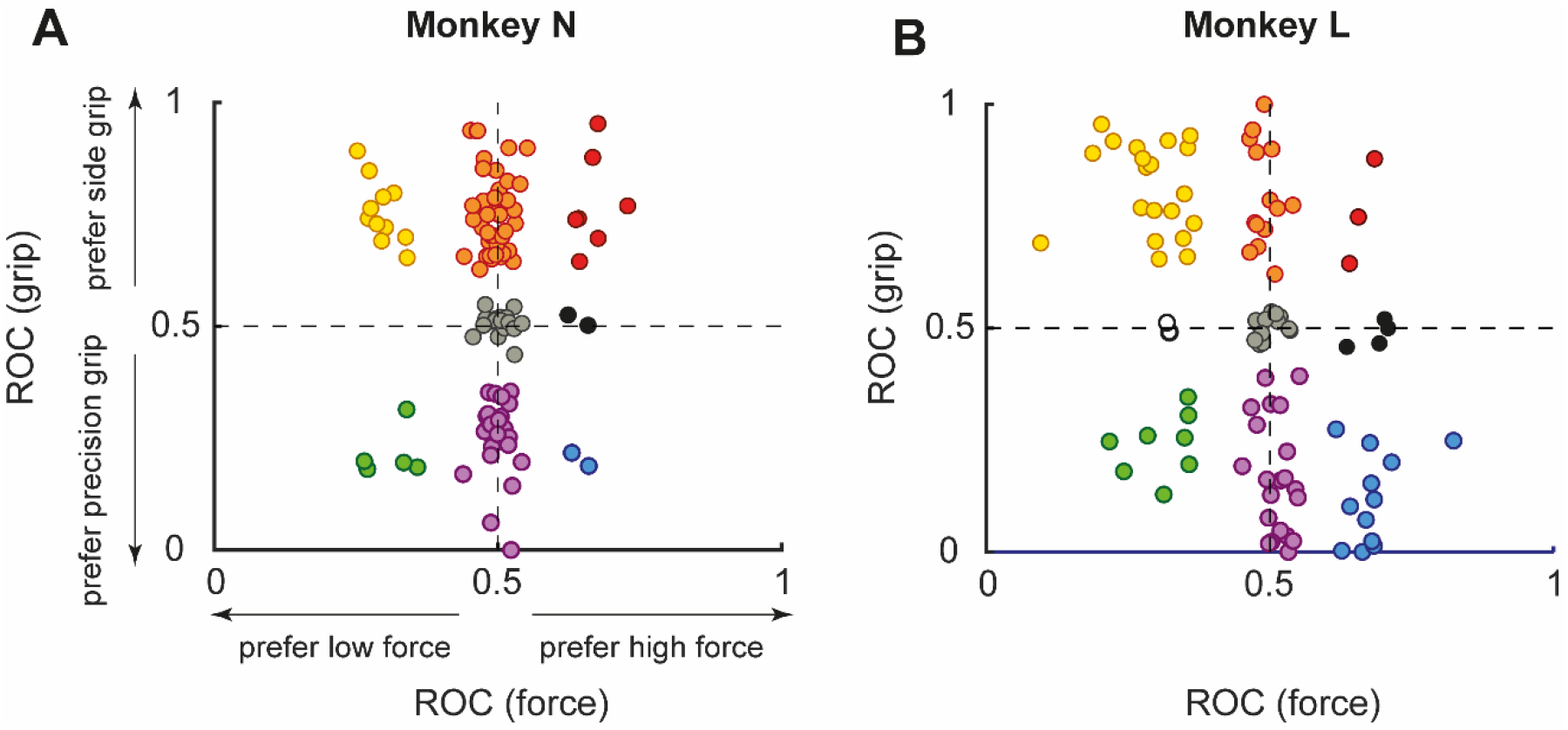
A heterogeneous firing rate code is used to represent grip and force in M1 neurons. A: Receiver-operating-characteristic index (ROC) obtained from the comparison of neural activity during side grip vs precision grip trials as a function of ROC comparing high force vs low force trials in monkey N. Each colored dot represents a neuron. Dot color indicates the cluster from which the neuron belongs to. B: Same as in A, for monkey L.

Consistent with the mutual information analysis, we observed a few neurons purely modulated by force level (2/110 in monkey N; black dots) and that most of the force representations came from neurons that were also modulated by grip. Interestingly, high force could be represented by either an increase (11/110 neurons) or a decrease (16/110 neurons) in the firing rate. Regarding grip type, it could be represented in two manners: either by only grip selective neurons (65/110) or by grip and force modulated (25/110) neurons. Among all of these, 61 units showed an increase while 29 decreased their firing rate during the side grip condition.

Figure 5B shows that these results are also observed in monkey L, namely, that force representations were embedded in neurons that were also modulated by grip type (43/97). Again, we found a heterogeneous neural force code, with some neurons increasing their firing rate in response to high force trials (17/97) and others decreasing their activity during the same condition (30/97). Less than 20% of all neurons were not modulated by either grip type or force level (16% or 18/110 for monkey N and 13% or 13/97 for monkey L). Thus, the mutual information and ROC analyses consistently demonstrate that grip type engaged a large population of grip specific neurons and that, conversely, force level was represented by smaller number of neurons that, in addition to force also contained grip information.

### Population encoding of grip type and force level

We performed a population analysis using demixed principal components (dPCs) which estimated the percentage of variance associated with (1) condition-independent modulations (related to the temporal structure of the task), (2) grip type, and (3) force level Figure 6A shows the normalized firing rates of all M1 neurons projected onto dPC #1. We found that the largest proportion of population variance is explained by condition independent activity that is related to the temporal structure of the task, i.e. to the moment at which the monkeys initiate and release the grasp on the object.

**Figure 6.**
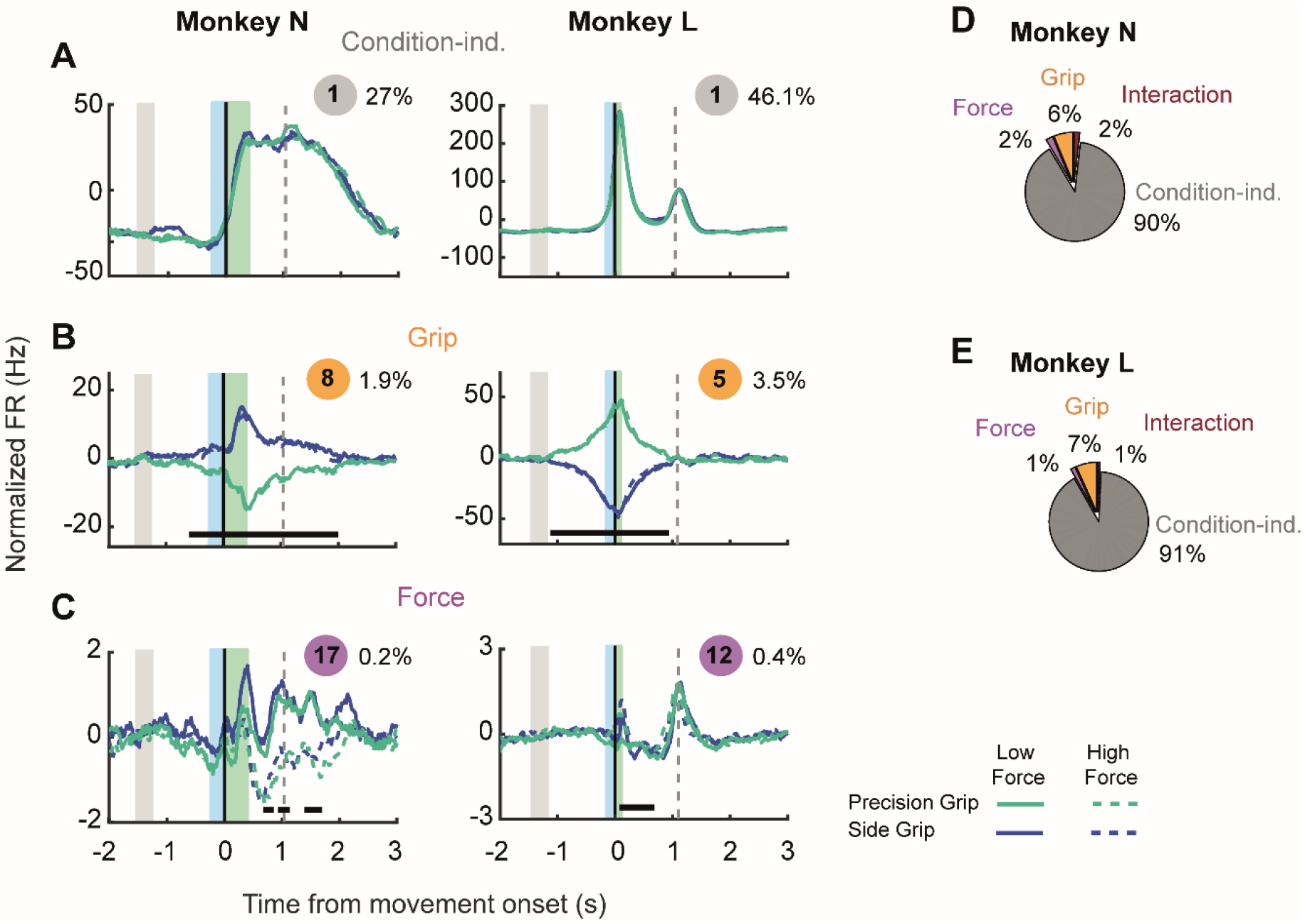
Population-level representations of grip type and force level in M1 using dPCA. Firing rates were projected onto the demixed principal component (dPC) that explains the most variance associated with each task parameter (condition-independent, grip and force). A: First conditionindependent component (component #1) for monkey N (left) and monkey L (right). B: First significant grip component for monkey N (component #8, left) and monkey L (component #5, right). C: First significant component associated with force for monkey N (component #17, left) and monkey L (component #12, right). Activity was aligned to movement onset (vertical black solid line). Grey shading indicates the presence of the grip cue. Blue and green shading indicate mean reaction and movement times respectively. Vertical dashed line corresponds to the average time of object release. Component number and percentage of explained variance are shown at the top right corner of each panel. Black solid line at the bottom indicates time points at which the task parameter could be decoded from the neural activity at single-trial level using its associated dPC as a classifier. D: Percentages of total explained variance by each task parameter for monkey N. E: Same as in D, for monkey L.

It is interesting to note that dPC #1 in monkey N showed a less pronounced slope in the increase of activity associated with movement onset, and this might be related to the slower reaction and movement times of the animal in comparison of monkey L. This increase was held during the entire movement period, with activity decreasing after object release. Conversely, monkey L temporal dynamic was overall faster, with a more pronounced slope during the peri-movement period and a sudden drop after object touch. This was followed by a small increase in activity in response to object release.

The dynamics of the dPCs related to grip type are shown in Figure 6B. It must be noted that the amount of variance associated with grip type are an order of magnitude smaller as compared to those related to the temporal structure of the task. In the case of monkey N, dPC #8 explained 1.9% of total variance, but this was enough to decode hand grip before movement onset. For monkey L, dPC #5 explained 3.5% of total variance and allowed to predict the monkey’s hand grip hundreds of milliseconds before the animal received the go cue (much earlier compared to monkey N). This neural representation was maintained until object release and was not held any further, as opposed to what we observed in monkey N.

The dynamics of the activity related to force level are shown in Figure 6C. In monkey N, dPC #17 explained 0.2% of total variance, and it allowed to decode force level later in the trials and for shorter windows as compared to grip type. Similarly to what we observed in grip decoding, force population signal was maintained even after object release in monkey N. For monkey L, dPC #12 allowed to predict force trials in a more consistent way, starting after movement onset and finishing before object release. Figure 6D and 6E show the total explained variance for each task variable in monkey N and monkey L, respectively. In both cases, condition-independent components explained about 90% of total neural variance. This was followed by grip that explained 6% and 7% of total variance in monkey N and L, respectively. Force explained only 2% of variance in the case of monkey N and 1% for monkey L. Finally, the interaction of grip and force explained less than 2% of total variance and none of the interaction dPCs were significant.

### Population activity allows decoding grip type and grip-force during single trials

Given that the neurons were simultaneously recorded, we used SVM classifiers to test whether population activity allowed the decoding of grip type and force during single trials. Figures 7A and 8A show the decoding accuracy of the grip type and grip-force classifiers as a function of time in monkeys N and L, respectively. For both animals, above-chance decoding for grip type started almost immediately after the onset of the grip cue and continued during movement execution. In contrast, significant decoding of force level was present only during movement execution and its magnitude was smaller than for grip. Next, to assess how stable were these neural representations throughout time, we performed a cross-temporal decoding analysis. Specifically, we trained and tested both classifiers using all possible pairs of time bins during the peri-movement period of the task. This method produces a classification accuracy matrix where the off-diagonal values are a measure of how generalizable the neural representation of grip type or force at a particular time bin is, when directly compared with the corresponding diagonal values, where the training (x-axis) and testing (y-axis) of the classifier are over the same time bin. Hence, the level of generalization across time bins tells us how robustly a set of neurons encode grip type or force across time during task performance. Figures 7B and 8B show the heatmaps of the cross-temporal decoding accuracies for grip and force classifiers using neural activity from monkeys N and L, respectively. For monkey L, grip decoding was more stable in time (static), with two generalization clusters, one during movement preparation, and another during grip execution. This is more evident in the binary matrix of Figure 8D, where the significant static off-decoding bins are depicted in yellow.

**Figure 7.**
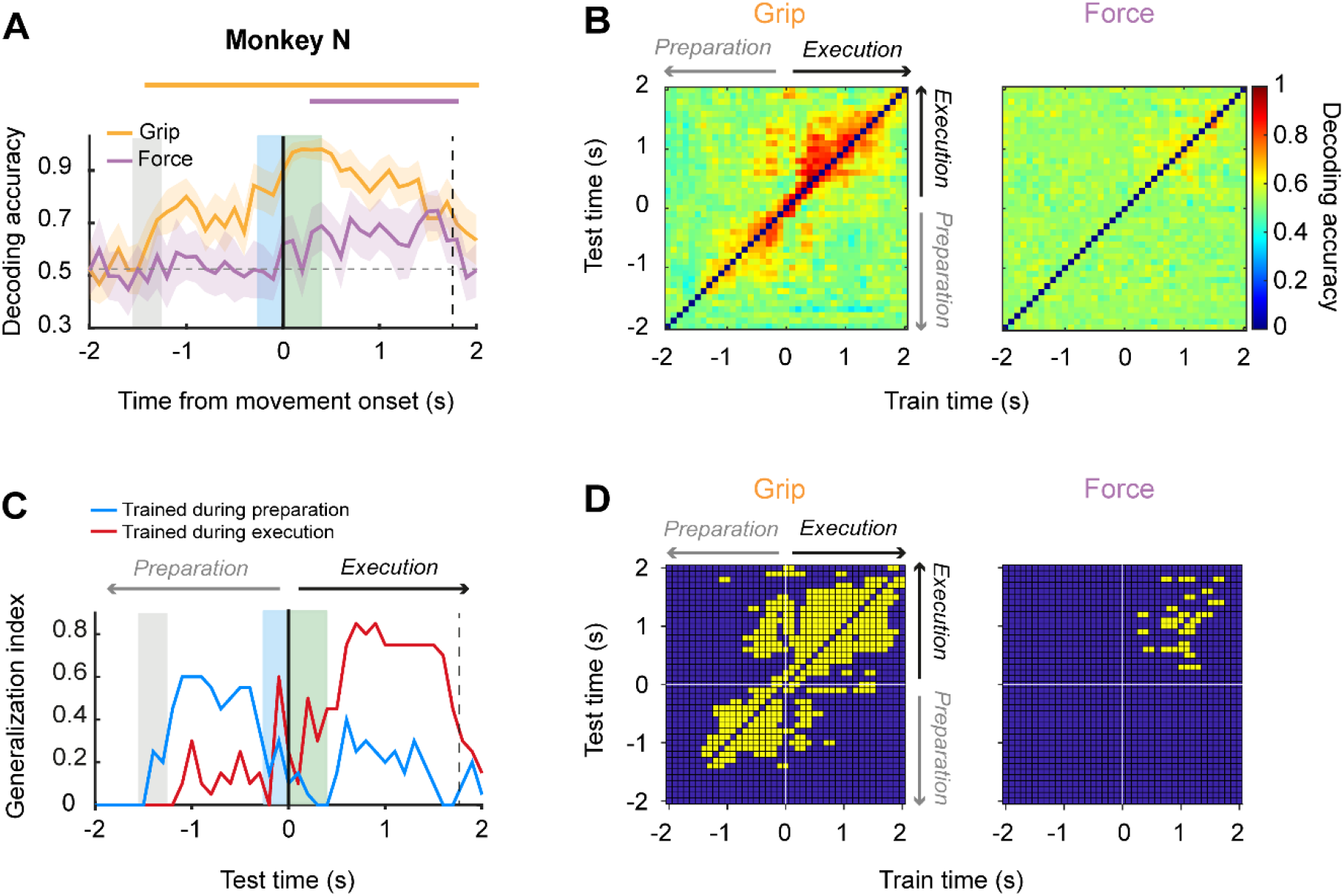
Temporal stability of grip and force neural representations in M1 (monkey N). A: Decoding accuracy of grip type (yellow solid line) and force level (purple solid line) as a function of time using an SVM classifier. Solid lines at the top indicate the times at which accuracy was significantly above chance (p < 0.01). Neural activity was aligned to the movement onset, indicated by the black vertical line. Gray shading indicates display of the grip cue. Blue and green shading indicate mean reaction and movement times, respectively. Vertical dashed line corresponds to average time of object release. B: Cross-temporal decoding of grip (left) and force (right) during the peri-movement period of the task. C: Generalization index estimated as the proportion of static bins resulting from the cross-temporal decoding of grip as a function of time when training and testing the classifier during preparatory and executive periods of the task. D: Stable time points of the crosstemporal decoding of grip (left) and force (right). Yellow squares indicate significantly stable time bins.

**Figure 8.**
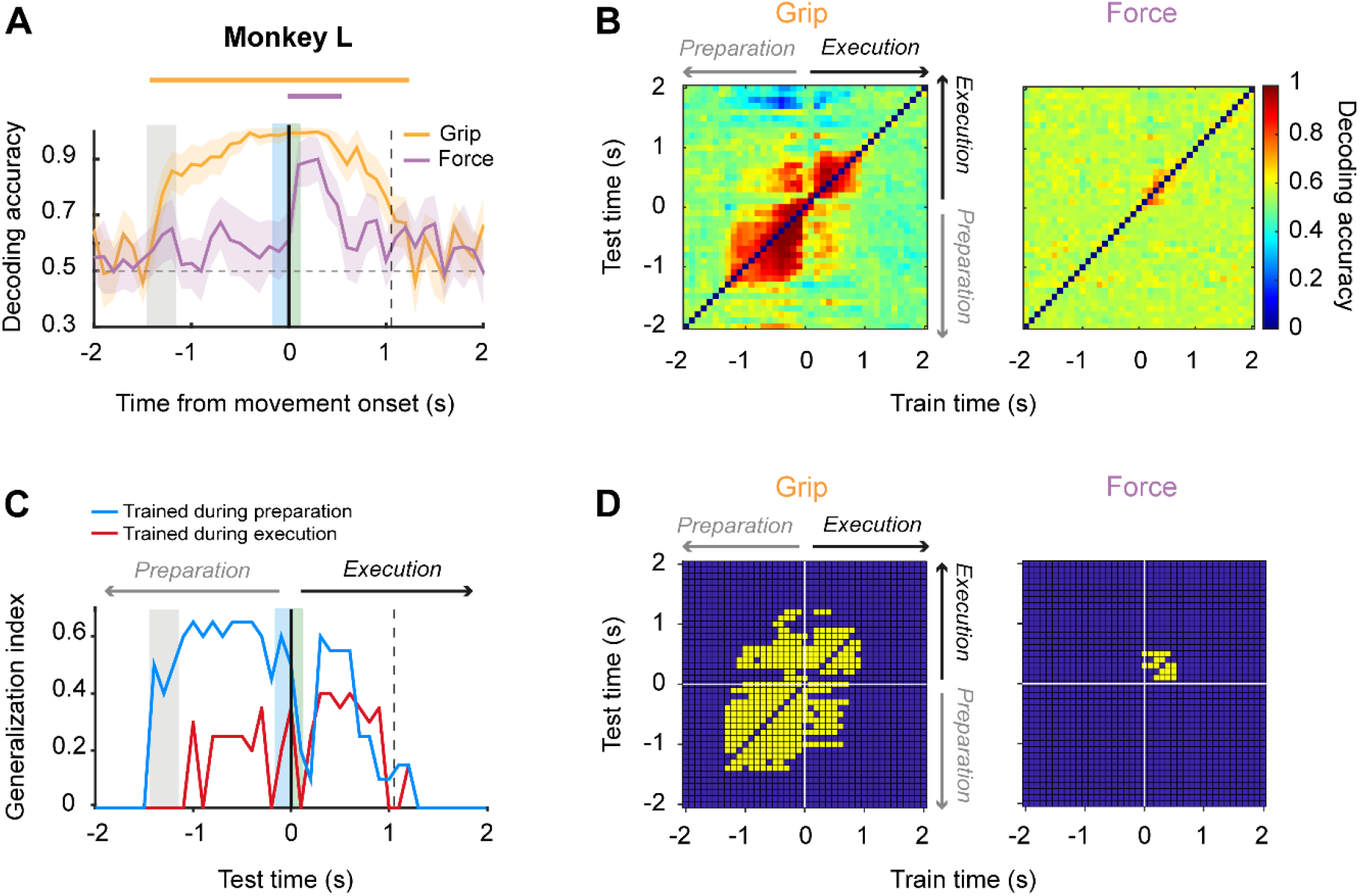
Temporal stability of grip and force neural representations in M1 (monkey L). A: Decoding accuracy of grip type (yellow solid line) and force level (purple solid line) as a function of time using an SVM classifier. Solid lines at the top indicate the times at which accuracy was significantly above chance (p < 0.01). Neural activity was aligned to the movement onset, indicated by the black vertical line. Gray shading indicates display of the grip cue. Blue and green shading indicate mean reaction and movement times, respectively. Vertical dashed line corresponds to average time of object release. B: Cross-temporal decoding of grip (left) and force (right) during the peri-movement period of the task. C: Generalization index estimated as the proportion of static bins resulting from the cross-temporal decoding of grip as a function of time when training and testing the classifier during preparatory and executive periods of the task. D: Stable time points of the cross-temporal decoding of grip (left) and force (right). Yellow squares indicate significantly stable time bins.

In contrast to grip type, force decoding was temporally less stable (dynamic), with only a few significant off-diagonal time bins, all of them close to the on-diagonal time bins, and that occurred only during the movement execution (Figure 8D). In the case of monkey N, grip type decoding was also more temporally stable than force (Figure 7B). However, stable time bins were distributed more sparsely across preparation and execution periods, with more static bins clustered within the execution period of the task.

We developed a generalization index that corresponds to the proportion of SVM bins that were classified as static when cross-temporally decoding grip and force across the preparation and execution epochs of the task. Hence, these indexes summarize how the significant decoding during the preparation epoch generalizes within the preparation and for the execution, or how the significant decoding during execution generalizes for the preparation and execution epochs. Figures 7C and 8C show the generalization index as a function of test time when the classifier was trained during preparation (blue line) and execution (red) for monkeys N and L, respectively. It is evident that there is a generalization of grip type decoding from preparation to execution and vice versa, suggesting that the motor cortical populations possessed information about the type of grip across task epochs, although there is a clear bias for a larger number of static bins when training and testing the classifier in the same epoch. Interestingly, grip type generalization was more pronounced during preparation for monkey N, whereas decoding during execution generalized more in monkey L.

## Discussion

In this study we aimed to determine how M1 neurons are involved in a reach-to-grasp behavioral task in monkeys. Our findings support three conclusions. First, we found that at the single-unit and population levels grip type had a large effect on motor cortical activity, while that force level explained less variance of the neural responses. Second, these parameters were encoded serially and independently from each other, even when the engaged neural populations were partially overlapping. Third, the neural signals linked to grip type emerged right after grip instruction and were maintained after movement onset, forming a long lasting and stable signal for precision or side grip during the preparation and execution epochs of the task. Thus, the categorical grip signal was more static compared to force representations, which occurred later and were more phasic. Our findings support the notion that M1 is strongly engaged in encoding grip type, with neural populations distinguishing between the two grips across the preparation and execution epochs of the task. In contrast, force was encoded phasically during movement execution and was mostly independent of grip type.

Both animals exhibited considerable differences in the timing of their actions, and neural activity followed a similar temporal pattern as behavior. Thus, the mean firing rate correlated with movement execution, with most neurons firing at the time of contact with the object across animals. Monkey L performed the task faster, and therefore neural responses peaked earlier compared to monkey N. We expected to find neural modulations around movement execution, since M1 activation during voluntary movement has been extensively documented in the literature (Cheney & Fetz, 1980; Evarts, 1968; Georgopoulos et al., 2007; Kakei et al., 1999; Merchant et al., 2004; Muir & Lemon, 1983; Naselaris et al., 2006; Tanji & Evarts, 1976). However, here we were interested in determining whether modulations in the firing rate were related to relevant movement parameters to solve the task: grip type and force level. Moreover, we aimed to determine whether these parameters could be represented as abstract categories that correlated with behavior.

### Grip type is represented at both single-unit and population levels in M1

For both animals, most neurons carried a significant amount of grip information. Neural representations of grip emerged during the delay period and peaked near movement onset. Since monkeys had to perform either a side or a precision grip, it is important to highlight the differences between them, especially in the motor coordination required for each. While a side grip demands the coordination of the thumb and the rest of the fingers as a group, a precision grip is a finer, and therefore a more controlled movement, that requires independent coordination of the fingers to grasp an object using only the thumb and the index finger (Almécija & Sherwood, 2016; Edin et al., 1992; Jeannerod et al., 1995). Previous studies have shown that M1 neurons fire more during a precision grip compared to a power grip. This neural modulation does not seem to represent muscle engagement exclusively, since there is not an absolute correlation between net force and the firing rate of individual neurons of the pyramidal tract in M1 (Maier et al., 1993; Muir & Lemon, 1983). A possibility is that action complexity and therefore, the level of motor control needed, could modulate neural activity, and send specific motor commands to the spinal cord related to different hand grips (Muir & Lemon, 1983). Whether neural activity in M1 represents muscles or movements has been a topic of debate for years. Evidence supports the notion that both can be equally represented in M1 neural activity (Kakei et al., 1999; Wang et al., 2022). Given the limitations of our study, we cannot fully discern whether anticipatory coding of grip reflects differential muscle activations or grip type as an abstract category, since we did not perform electromyography (EMG) nor kinematic analysis. However, our findings may support the latter view. We found a heterogeneous firing rate code for representing grip types in M1 with some neurons firing more during a precision grip, while others “preferred” a side grip. This indicates that grip type in M1 is encoded independently of the pattern of muscle activation required for performing either grip type, suggesting that M1 seems to represent grip categorically. Moreover, we also found a negative correlation between the amount of anticipatory grip information and reaction and movement times when comparing both animals, with the faster animal also exhibiting more preparatory grip information. This could imply that the categorical representation of grip in M1 may be actively used by the brain to achieve proper motor, as well as temporal precision.

Grip type was also represented at the neural population level. Analysis of population activity using dPCA demonstrated that, in both animals, representation of grip emerges during the delay period, while the animal prepares its movement. Similar to what we found at the single-unit level, significant decoding of grip was possible earlier in monkey L, almost immediately after the grip cue. In addition, our findings at the neural population-level are consistent with what has been previously reported in the literature for M1 and other grasping-related areas like the anterior intraparietal area (AIP) and the anterior portion of the ventral premotor cortex (area F5), in which time is the most represented task parameter, followed by grip and lastly, force (Intveld et al., 2018; Rastogi et al., 2021).

### Force level is less represented than grip in single units and the neural population of M1

One of the main advantages of this task design is that it allowed us to dissociate and study how grip type and force level are represented in the neural activity. This was possible because both variables were separated in time, given that monkeys first had to grasp the object and then pull it. The cues for each variable were also temporally apart, allowing us to study their preparation independently. While the animals had a 1-second delay to prepare their hand grip, pulling force had to be prepared “on the go”, therefore, force preparation overlapped with movement execution. Overall, we found that force was less represented than grip in both animals. At the single-unit level, we found less force-related neurons and also, they carried less force information, compared to grip-related neurons. Most of these cells were co-modulated by grip, implying that coding of force in M1 mostly occurs through mixed-selective neural subpopulations (Kobak et al., 2016; Rigotti et al., 2013). At the population level we found that force explained less neural variance than grip, consistent with what we found in single units.

At first, we considered that a possible explanation for the differences between grip and force coding could reside on the time the animals had to prepare both. We thought of the possibility that having less time to prepare the appropriate force level could impact on the amount of explained variance within the neural population of M1. However, a previous study using dPCA to identify population representations of both parameters in grasping related areas, including M1, also showed that force explained less neural variance than grip, even when both grip and force were simultaneously planned within the same time window (Intveld et al., 2018).

Remarkably, all force and grip representing subpopulations showed very diverse neural responses, with neurons firing more during a certain grip type and/or force level but with no dependency between both variables. Furthermore, as what we found for grip coding, force-representing neurons could preferentially increase their firing rate during the high force or low force conditions, suggesting that this activation does not depend, at least linearly, on muscle engagement.

### Grip and force are represented independently in M1 at both single-unit and population level

Several studies have found that motor-related areas like the posterior parietal cortex (PPC), AIP, the dorsal and ventral premotor cortices (PMd and PMv, respectively) and M1 actively participate in grip and force control during grasping (Chib et al., 2009; Davare et al., 2006; Intveld et al., 2018). However, it is not clear whether these crucial aspects of grasping behavior are a result of the same neural process, or they are driven by separate neural mechanisms. Whether M1 represents muscle activity, movement parameters or generates neural states that facilitate movement remains under discussion. Evidence suggests that it is likely all these features are encoded in different single-unit and population-level neural codes (Cheney et al., 1980; Churchland et al., 2012; Evarts, 1968; Georgopoulos et al., 1986; Taira et al., 1996; Tanji & Evarts, 1976). While some studies have pointed a common neural substrate for grip and force control in M1 (Hepp-Reymond et al., 1999; Maier et al., 1993), more recent findings suggest that these parameters are represented independently at the single-unit level (Hendrix et al., 2009). Our findings support the latter notion and extend it to the neural population level, with none of the grip-force interaction dPCs resulting significant, supporting the view that both parameters are categorically and independently encoded in M1 neural activity.

### Grip representations are more stable than force in M1

Lastly, we evaluated the temporal stability of grip and force neural representations. We found that grip coding was more stable in time (hence, static), and that it was maintained by the neural population throughout the delay period, during movement preparation. On the other hand, coding of force was more dynamic, and was active mainly during movement execution. The neural mechanisms behind the static coding scheme are still being studied. However, it has been proposed that neural representations are sustained in time by reverberating connections between pyramidal neurons through activation of NMDA receptors (Wang et al., 2013).To further explore the static coding of grip in M1, we developed a generalization index as a measure of the temporal generalization of grip encoding across preparatory and executive periods of the task by the same neural population (Crowe et al., 2014; Gámez et al., 2019). We found that grip encoding generalizes across both periods of the task despite the temporal differences in movement timing between animals. Therefore, our results indicate that M1 neural populations can keep a sustained representation of grip type during preparation and execution of movement. Interestingly, we did not observe this during force coding, suggesting that the representation of task-relevant movement parameters in M1 can be embedded in distinct coding schemes that change flexibly, according to task demands. Previous studies have shown that static and dynamic coding schemes can coexist within the same cortical area (i.e. the prefrontal cortex) and it is possible to switch from one to another several times according to factors like task requirements and learning (Ceccarelli et al., 2023).

In summary, we found that grip type and force level are represented independently by M1 neural activity during grasping at both single-unit and population levels. Encoding of grip type was carried by both mixed-selective and grip-selective neurons, while force was represented by a smaller group of mixed-selective neurons. These representations do not seem to be associated to stereotyped muscle activation patterns of each movement, suggesting that they may be abstract and are informative about future behavior. In both animals, grip was more represented than force and neural representations of grip were more static engaging similar neural populations across task epochs, while force representations were short lived. This indicates that M1, like other associative areas such as the prefrontal cortex (Ceccarelli et al., 2023; Merchant et al., 2011) can switch from a static to a dynamic coding scheme in response to task demands. In addition, population-level neural representations allowed to predict grip and force independently at the singletrial level.

## Acknowledgements

This work was supported by Consejo Nacional de Humanidades, Ciencia y Tecnología (CONAHCYT) Grant: A1-S-8430, UNAM-DGAPA: IG200424, and UNAM-DGAPA-PASPA to H. Merchant, and by CONAHCYT: 319212 to V. de Lafuente. Adriana Moreno is a doctoral student from Programa de Doctorado en Ciencias Biomédicas, Universidad Nacional Autónoma de México (UNAM) and has received CONAHCYT fellowship 1003309. We thank Luis Prado, Raúl Paulín, María Antonieta Carbajo and Juan Ortiz for their technical assistance.

The authors declare no competing financial interests.

